# Pitch selectivity in ferret auditory cortex

**DOI:** 10.1101/2025.11.03.686220

**Authors:** Veronica Mae Tarka, Quentin Gaucher, Kerry Marie May Walker

**Affiliations:** Department of Physiology, Anatomy & Genetics, University of Oxford, Oxford, OX1 3PT, UK; Sorbonne Université, CNRS, Inserm, Institut de la Vision, F-75012 Paris, France

**Keywords:** Auditory cortex, Neural coding, sensory integration, temporal, spectral, pitch, perception

## Abstract

Our perception of pitch – the tonal quality of a sound at a fundamental frequency (F0) – is essential for musical melody, vocal communication, and attending to a voice in a crowded room. Pitch is distinct from simple frequency tuning. Although pitch perception can arise from either harmonic spectral structure or temporal regularity, it remains unclear how individual neurons represent these distinct cues. We recorded spiking activity in hundreds of ferret auditory cortical neurons while systematically varying harmonic and temporal pitch cues. A subset of neurons represented F0 based on harmonic content, another subset on temporal periodicity, and a third population exhibited invariant pitch tuning across both cue classes. These findings provide the first evidence for specialized pitch neurons in non-primates, and demonstrate that mammalian auditory cortex employs dual spectral–temporal mechanisms for extracting F0.

**Significance statement:** Pitch allows us to enjoy music, understand speech, and follow voices in noisy settings, yet how the brain represents pitch remains a long-standing question. Pitch selective neurons have been identified previously only in primates, limiting our ability to investigate the neural of pitch. Here, we show that ferret auditory cortex also contains such neurons, which extract pitch from both harmonic structure and temporal periodicity. This discovery reveals that the brain uses dual, complementary strategies for encoding pitch.

## Introduction

Sounds in nature are typically composed of many frequencies, yet we often perceive a natural sound to have a single tonal quality along a low-to-high scale, known as its pitch. Pitch perception supports recognition of musical melody, prosody in speech, and the segregation of sound sources in a complex acoustic environment, such as following a conversation in a noisy restuarant^1–4^. Despite its essential role in hearing, it remains unclear how pitch is extracted from acoustics and represented in the brain^5^.

Many sensory qualities are not represented by a labeled line code in the periphery. For example, color is a percept synthesized by the visual system that is not directly dictated by a single wavelength of light. Similarly, the perceived pitch is not simply the range of frequencies spanned by a sound. In the spectral domain, when all frequency components are integer multiples (i.e. harmonics) of a common fundamental frequency (F0), a pitch is perceived at the F0. This is true even for “missing fundamental” sounds, which contain no physical energy at F0 itself^6^. The frequency-tuned auditory nerve fibers of the cochlea can represent this spacing of harmonics as a place code using labeled lines, but only for lower “resolved” harmonics^7^. In the temporal domain, harmonic sounds are periodic, with a waveform repeating at 1/F0. Auditory nerve fibers tuned to higher, unresolved harmonics represent F0 in their spike timing by phase-locking to the sound envelope^8^. Psychophysical studies in humans and animals have shown that pitch can be perceived when sounds contain exclusively harmonic or temporal cues^9–12^. Therefore, the auditory system may use dual processing to identify F0.

Neurological studies^13–15^, lesion experiments in cats^16^, and neuroimaging in human listeners^13,17^ point to a key role of auditory cortex in pitch processing. Microelectrode studies across a range of mammalian species have described auditory cortical neurons sensitive to F0^18–20^, but only in marmosets have specialized “pitch neurons” been identified that extract F0 invariantly across sound types, including missing fundamentals^21^. These neurons were clustered at the low-frequency border of primary and secondary auditory cortex, raising the possibility of a specialized pitch area in the brain.

To investigate whether invariant representations of pitch exist in non-primate auditory cortex, we recorded the F0 tuning of individual neurons in ferret primary auditory cortex while presenting sounds that varied in their harmonic structure, phase, and temporal regularity. We found a population of pitch selective responses in auditory cortex, with some neurons integrating the labeled line-coded harmonic cues (“harmonicity neurons”) and others tuned to periodicity (“temporal neurons”). A subset of “pitch” neurons encoded F0 invariantly across both cue types, reflecting the unified perception of the pitch evoked by these different sounds. Importantly, the pure tone frequency tuning of these pitch selective neurons was not predictive of their F0-tuning, suggesting that cortical pitch representations necessarily integrate over multiple frequency components of a sound. These findings demonstrate that pitch selective neurons are not unique to primates.

## Results

We examined the hypothesis that pitch is represented in individual primary auditory cortical neurons of ferrets. We presented anaesthetized ferrets with a collection of 13 stimulus types that varied in harmonic components, phase, and the presence of a pink noise masker. Each stimulus type was presented across 17 different fundamental frequencies (1/4 octave increments across 250 – 4000 Hz). We recorded responses from 1266 well-isolated single neurons across 4 ferrets. This included 1077 neurons in A1 and 189 in AAF. We found that 872 (81%) of the single neurons we recorded were sound responsive (paired t-test, p<0.05). Of these sound-responsive neurons, 165 (19%) were sensitive to the F0 of periodic click trains (1-way ANOVA, p<0.05).

Because click trains contain both resolved and unresolved harmonics, we reasoned that neurons encoding F0 through either the harmonic or temporal envelope cues should be sensitive to the F0 of these click trains. Therefore, only sound-responsive neurons that were also sensitive to the F0 of click trains were included in subsequent analysis. We refer to this population as “F0-sensitive neurons”.

### Harmonicity neurons

We first aimed to test whether a subset of single neurons in the auditory cortex may extract pitch exclusively from resolved harmonics. We reasoned that such “harmonicity neurons” would derive a sound’s F0 by integrating the place code pattern of resolved harmonics across input neurons that are each tuned to individual harmonics of the sound. Neurons using such a strategy would be unable to extract pitch information from unresolved harmonics. They should therefore show similar F0-tuning across any sounds that contain resolved harmonics, and may be either unresponsive or untuned to the F0 of sounds containing only unresolved harmonics (Fig. 1a).

**Figure 1.**
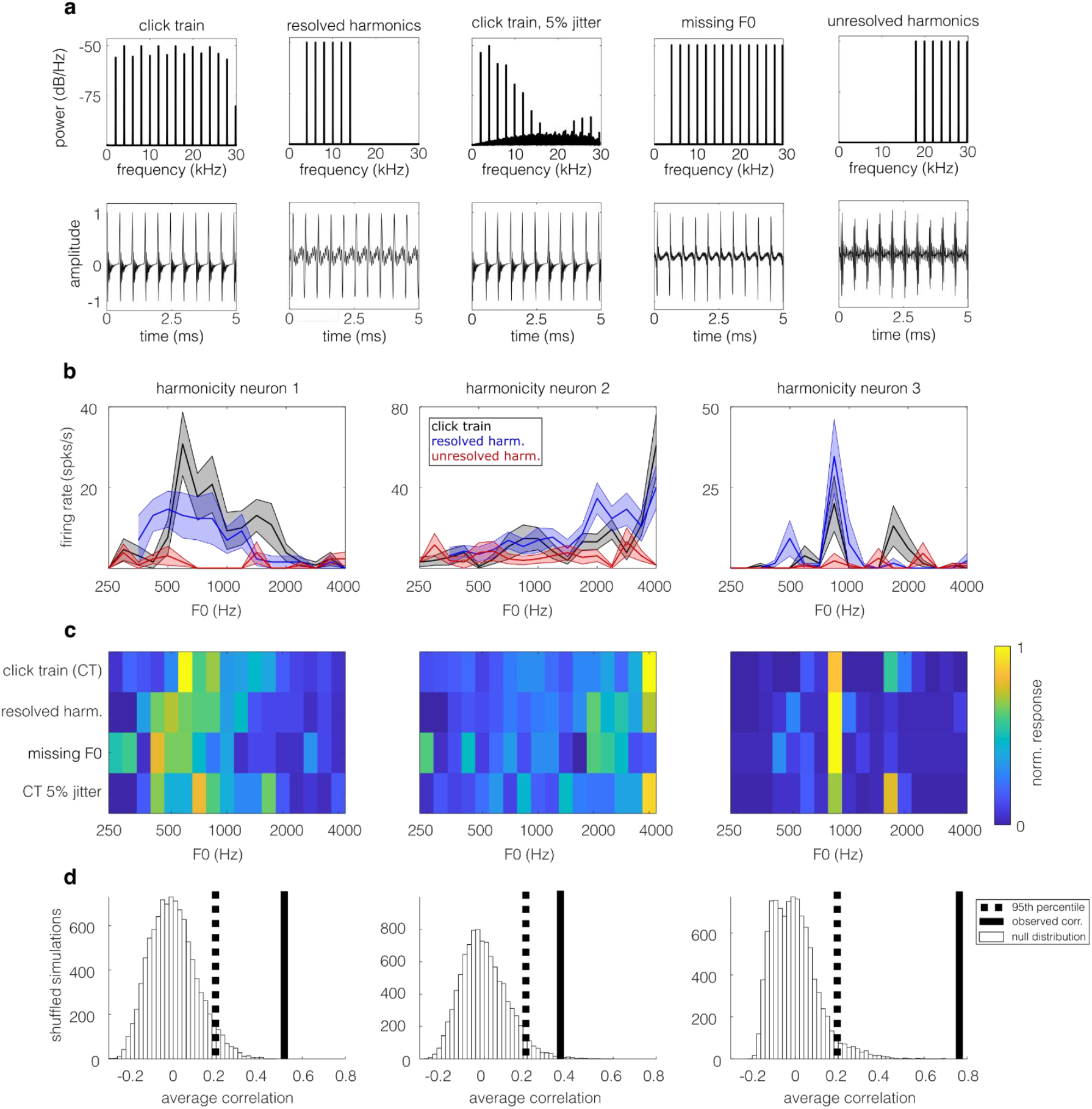
Harmonicity neurons derive F0 from resolved harmonics. **a**) Spectra (top row) and waveforms (bottom row) of the 5 different sound types used to classify harmonicity neurons. Each of these examples has an F0 of 2000 Hz. **b**) Each of the 3 panels shows the trial-averaged spike rate response of an example harmonicity neuron, as a function of F0. Shaded areas show the standard error of the mean (SEM). Harmonicity neurons derive F0 from resolved harmonics and therefore are sensitive to the F0 of click trains (black line) and resolved harmonics (blue line), but not to unresolved harmonics (red line). **c**) The color scale in each plot indicates the normalized trial-averaged spiking responses to 4 types of stimuli that contain resolved harmonics. **d**) In each plot, the solid black line indicates the average correlation between F0 tuning curves across all possible pairs of stimuli shown in **c.**The histogram (open bars) shows a null distribution of the average F0-tuning correlation calculated when the tuning curves in **c** were shuffled randomly across F0. The dashed black line shows the 95^th^ percentile of this null distribution. Results for the same 3 example harmonicity neurons are shown in **b, c**, and **d**.

Within the F0-sensitive neurons described above, we identified putative harmonicity neurons as those that were F0-sensitive to sound that contained only resolved harmonic complexes (1-way ANOVA, p<0.05), but were not F0-sensitive to tone complexes containing only higher, unresolved harmonics (1-way ANOVA, p≥0.05). This criterion identified 56 putative harmonicity neurons (34% of 165 F0-sensitive neurons). The F0 tuning curves of 3 putative harmonicity neurons are shown in Fig. 1b.

Note that this initial analysis identified neurons with spike rates that were modulated by the F0 of both click trains and resolved harmonic tones, but it does not require that the neuron show similar F0-tuning curves for these 2 stimuli. A harmonicity neuron should give a reliable read-out of a sound’s F0 across different types of sounds that contain resolved harmonics in order to support pitch perception. Therefore, we next tested if these putative harmonicity neurons showed consistent F0-tuning across sounds that contain resolved harmonics, but have different timbres. These example neurons maintained their tuning when we deleting all the energy at the frequency component corresponding to F0 (missing F0 sounds; Fig. 1c). Similarly, when we introduced temporal and spectral noise to the original click trains with 5% jitter in the timing of their individual clicks, these neurons were still tuned to the fundamental (CT 5% jitter; Fig. 1c). This invariant F0 tuning is visualized as vertical bands of high spike rates at similar F0s across all 4 resolved harmonic stimuli in Fig. 1c, although there is some variation in the width and peak of F0-tuning. Note that these neurons were not F0-sensitive when only temporal F0 cues were available (red lines in Fig. 1b), highlighting their dependence on resolved harmonics.

We used a bootstrapping approach to test if the timbre-invariant F0 tuning visualized in Fig. 1c was statistically significant. For each putative harmonicity neuron, we calculated the correlations between F0-tuning curves for all possible pairs of the 4 resolved harmonic sounds. We created a null distribution of these average correlations from F0-randomized versions of the same neuron’s responses. If the true F0-tuning curve correlation was larger than the 95^th^ percentile of the null correlation distribution, we concluded that the neuron showed significant generalization of pitch tuning across resolved harmonic stimuli (Fig. 1d). Of the 56 putative harmonicity neurons tested, 59% (n=33) showed significantly correlated F0-tuning across the resolved harmonic stimuli (p<0.05), and these were classified as harmonicity neurons.

### Temporal neurons

We next examined whether neurons in auditory cortex can extract F0 exclusively from unresolved harmonics. We hypothesized that such “temporal neurons” would derive a sound’s F0 by integrating the spike timings of inputs that are phase-locked to the sound’s temporal periodicity. Unlike harmonic neurons, they would show similar F0-tuning across any sounds that contain unresolved harmonics, regardless of whether resolved harmonics were also present (Fig. 2a). In principle, temporal periodicity could also be derived from the phase-locked activity of neurons responding to resolved harmonics, so temporal neurons may be either unresponsive, untuned, or tuned to the F0 of sounds containing only resolved harmonics.

**Figure 2.**
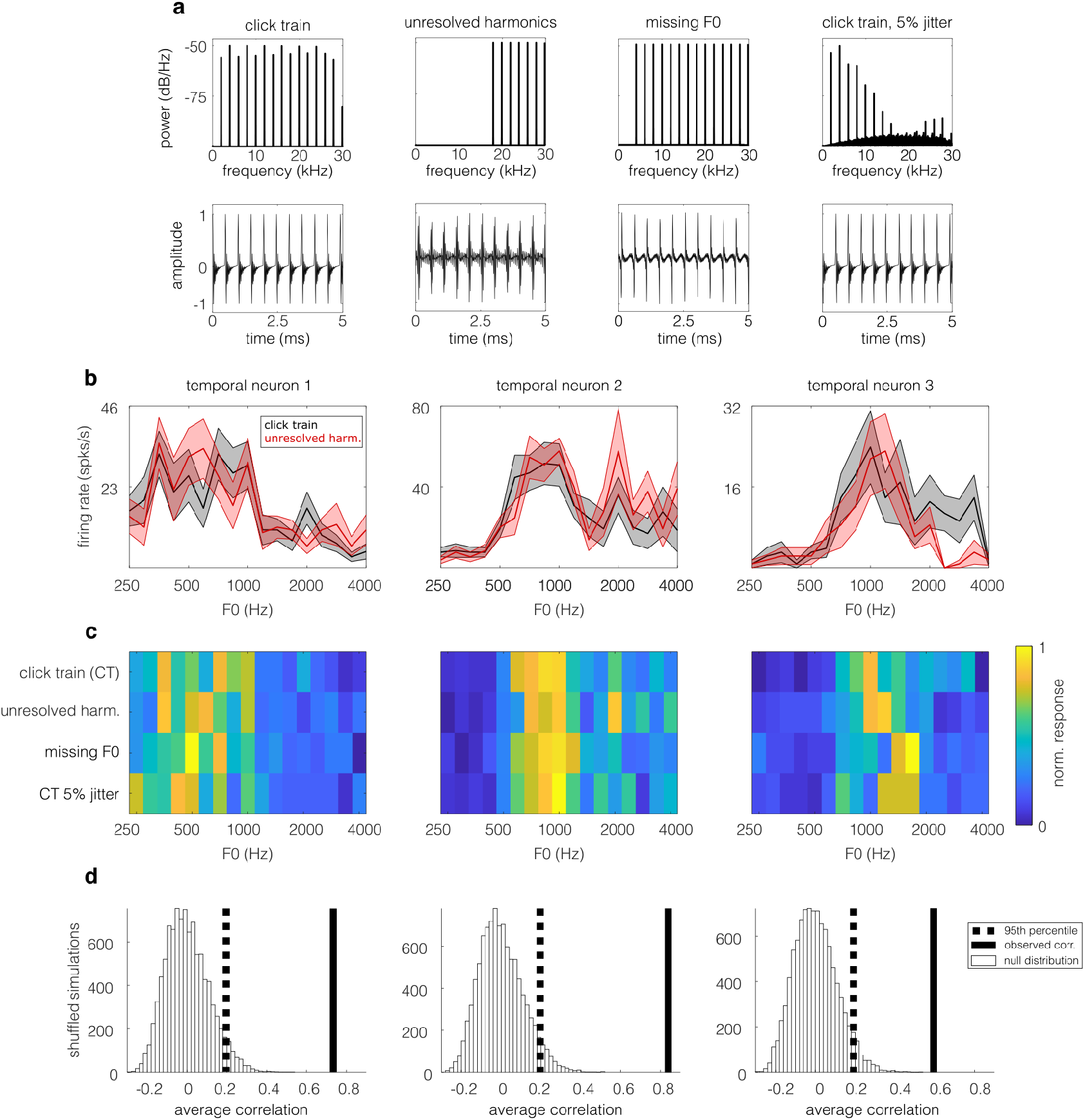
Temporal neurons derive F0 from unresolved harmonics. **a**) Spectra (top row) and waveforms (bottom row) of 4 different sound types used to classify temporal neurons (F0=2000 Hz). **b**) Each panel shows the trial-averaged spike rate response for an example temporal neuron as a function of F0. Shaded area shows the SEM. Temporal neurons show F0 sensitivity for click trains (black line) and unresolved harmonics (red line). **c**) The normalized trial-averaged responses to 4 types of stimuli that contain unresolved harmonics, plotted as in Fig. 1c. **d**) Results of bootstrapped correlation test of F0-tuning similarity across stimuli, plotted as in Fig. 1d. The same 3 example temporal neurons are shown in **b, c** and **d**.

We assessed periodicity tuning within our original set of 165 F0-sensitive neurons by identifying neurons that were sensitive to the F0 (1-way ANOVA, p<0.05) of click trains that contained either: (1) a combination of resolved and unresolved harmonics (Fig 2a; click train), or only high, unresolved harmonics (Fig 2a; high harmonics). The F0 tuning curves of 3 neurons reaching these two criteria are shown in Fig. 2b. In total, 44 neurons (27% of 165 F0-sensitive neurons) showed this temporal-based F0 sensitivity.

As with harmonicity neurons, these putative temporal neurons should provide a reliable read-out of a sound’s F0 across stimuli with different frequency spectra in order to support pitch perception. We therefore tested if these putative temporal neurons showed consistent F0-tuning across sounds that contain unresolved harmonics but have different frequency spectra (Fig. 2a). These comprised the click trains and high harmonic stimuli already described, as well as missing fundamental sounds and click trains with 5% jitter. A visual inspection of the spike rate responses of the 3 example putative temporal neurons showed a striking similarity in F0-tuning across the 4 sound types (Fig. 2c). We applied the same bootstrapping procedure described above for harmonicity neurons (Fig. 1d), and found that over 84% of these putative temporal neurons showed statistically similar F0-tuning across the 4 stimuli (p<0.05; Fig. 2d). Based on their ability to encode the F0 of unresolved harmonics across a wide variety of sound spectra, we classified these as temporal neurons (n=37).

### Temporal neurons are sensitive to phase manipulations

If temporal neurons primarily derive F0 from the sound’s periodicity, they should be sensitive to manipulations of the sound’s temporal fine structure. In the stimuli considered thus far, all harmonics were presented in phase. Here, we compare the responses of temporal neurons to 2 versions of the high harmonic stimuli in which the temporal cues were disrupted through phase manipulations. In the alternating phase (“alt. phase”) condition, the starting phase of consecutive harmonics alternated between sine and cosine phase. This creates a temporal periodicity at F0 and a second, comparatively weaker periodicity at 2F0 (Fig. 3a, middle). We therefore predicted that the tuning curves of temporal neurons might be shifted to a lower F0 in response to the alternating phase stimuli compared to the same sound presented in phase. In the randomized phase condition (“rand. phase”), the phase of each harmonic was individually randomized, in order to diminish the temporal periodicity at F0 and create a flatter temporal envelope (Fig. 3a, right). We predicted that temporal neurons would show poor F0-tuning in response to the randomized phase stimulus compared to the same harmonics presented in phase. It is important to note that both these manipulations left the harmonic content of the stimuli as revealed in the sounds’ spectra completely unchanged (Fig. 3a), so any changes in neuronal response would be due entirely to temporal sensitivity.

**Figure 3.**
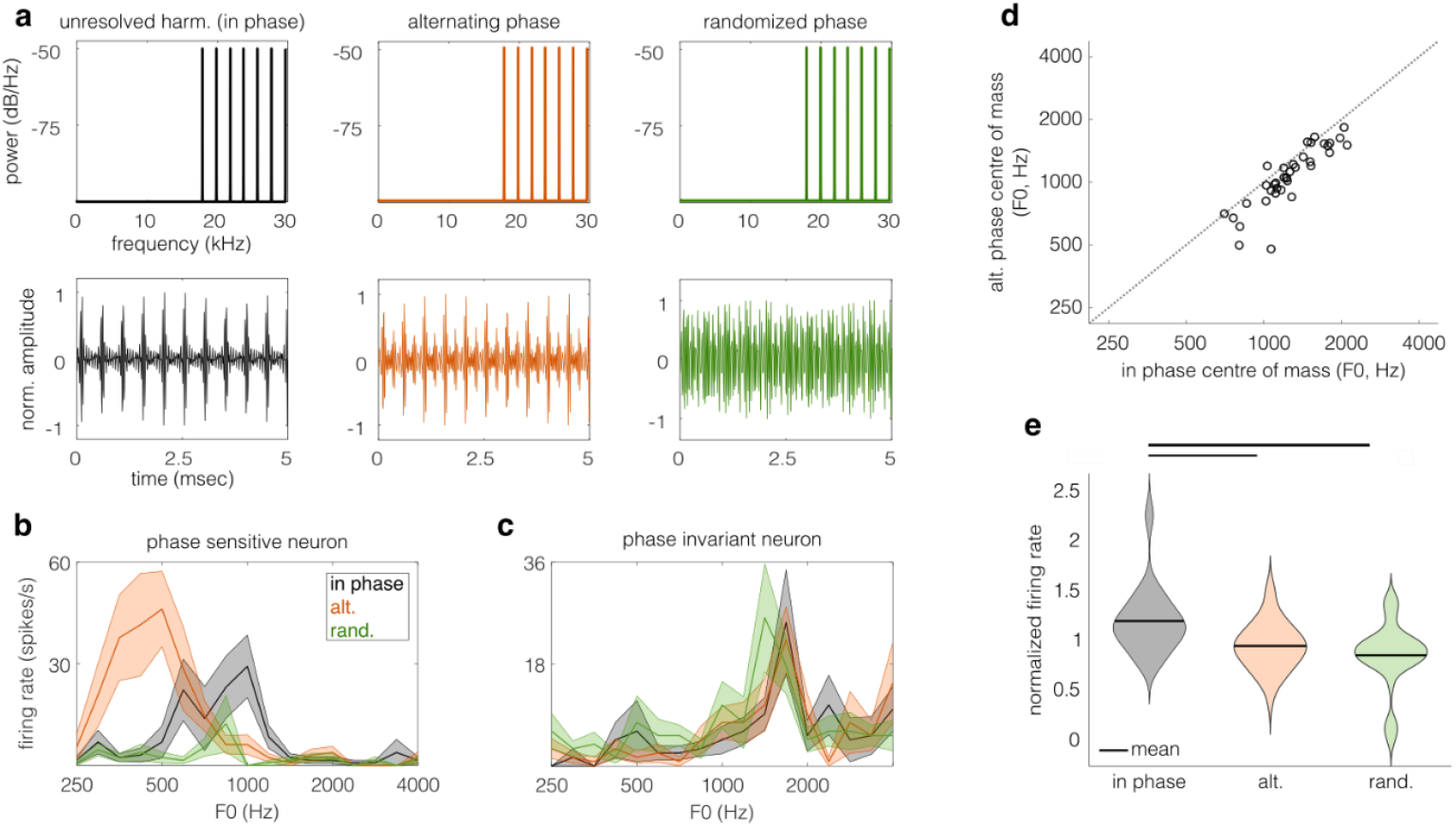
Temporal neurons are sensitive to phase manipulations. **a**) Spectra (top) and temporal waveforms (bottom) of stimuli in the 3 phase conditions are shown. The left panels show a tone complex containing high, unresolved harmonics, with each harmonic component presented in cosine phase (black). This stimulus has a strong temporal periodicity at F0. The middle panels show the same high harmonic complex presented with “alternating phase” (orange). The right panels show the high harmonic stimulus, with each harmonic component starting in randomized phase (green). **b**) F0-tuning of a phase-sensitive temporal neuron. The firing rate during each of the 3 high harmonic stimuli is shown as a function of F0. **c**) F0-tuning of a phase-invariant neuron, plotted as in **b. d**) The center of mass of a neuron’s F0-tuning curve was defined as the weighted average of the trial-averaged responses elicited across all F0s presented. The scatterplot shows that most temporal neurons had a lower center of mass for alternating phase stimuli compared to the same stimuli presented in phase (paired t-test; p<0.05). **e**) Each temporal neuron’s response to a given phase manipulation was taken as the average of its response at the best F0 (+/-0.25 octaves) of the in phase F0-tuning curve. This response was then normalized across all phase manipulations. Neural response magnitude was smaller for the alternating and random phase stimuli than the in phase stimuli (1-way ANOVA with Tukey-Kramer HSD test; thin line: p<0.01, thick line: p<0.001).

We found that a subset of temporal neurons was sensitive to phase manipulations. An example of a phase-sensitive temporal neuron is shown in Fig. 3b. This neuron was tuned to a lower F0 for the alternating phase stimulus (orange line) compared to the in-phase version of this sound (black line), reflecting the increased periodicity of the alternating phase version. Furthermore, this neuron showed weaker responses to sounds with randomized phase, even when they were presented at the preferred F0 (green line). Therefore, this example neuron showed the phase sensitivity predicted for temporal neurons. Not all temporal neurons showed this level of phase sensitivity. A second example neuron shown (Fig. 3c) is one of the most phase-invariant temporal neurons in our dataset, showing similar F0-tuning across all three high harmonic stimuli.

Phase sensitivity was found to be a common feature of temporal neurons overall. Their F0 tuning was consistently and significantly lower in response to alternating phase stimuli than to the same harmonics presented in phase (one-tailed paired t-test; t(36)=6.1, p=2.9×10^-7^; Fig. 3d). These phase manipulations also elicited weaker spiking responses compared to same harmonics presented in phase (1-way ANOVA, F(2)=12.86, p=9.79×10^-6^; Fig. 3e). Temporal neurons produced weaker spike rate responses at their preferred F0 in response to alternating phase stimuli (Tukey-Kramer HSD test; p=0.0015) and random phase stimuli (Tukey-Kramer HSD test; p=1.00×10^-5^), compared to harmonics in phase. Therefore, the population of temporal neurons is sensitive to fine temporal manipulations, even across sounds with identical spectral content.

### Pitch neurons encode F0 using both harmonic and temporal cues

Having identified harmonicity neurons that derive F0 from resolved harmonics and temporal neurons that derive F0 from the periodicity of unresolved harmonics, we next investigated whether there may exist more general “pitch neurons” that can derive F0 from both these acoustic cues. To identify pitch neurons, we applied a protocol similar to the one used for temporal and harmonicity neurons. First, we defined putative pitch neurons as those that were F0-sensitive for click trains, resolved harmonics, and unresolved harmonics (Fig. 4a), combining the inclusion criteria of harmonicity and temporal neurons. We found 32 neurons that passed this criterion, and three examples are shown in Fig. 4b. To determine if these putative pitch neurons could generalize their F0-tuning across a wide range of sounds, we used the same bootstrapping approach described above for harmonicity and temporal neurons to test if F0-tuning was similar across 5 stimulus classes: click trains, low harmonics, high harmonics, missing F0 sounds, and click trains with modest (5%) jitter (Fig. 4a). While these 5 stimuli have widely different frequency spectra and temporal fine structure, they should all evoke a percept of pitch in both humans and ferrets^12^.

**Figure 4.**
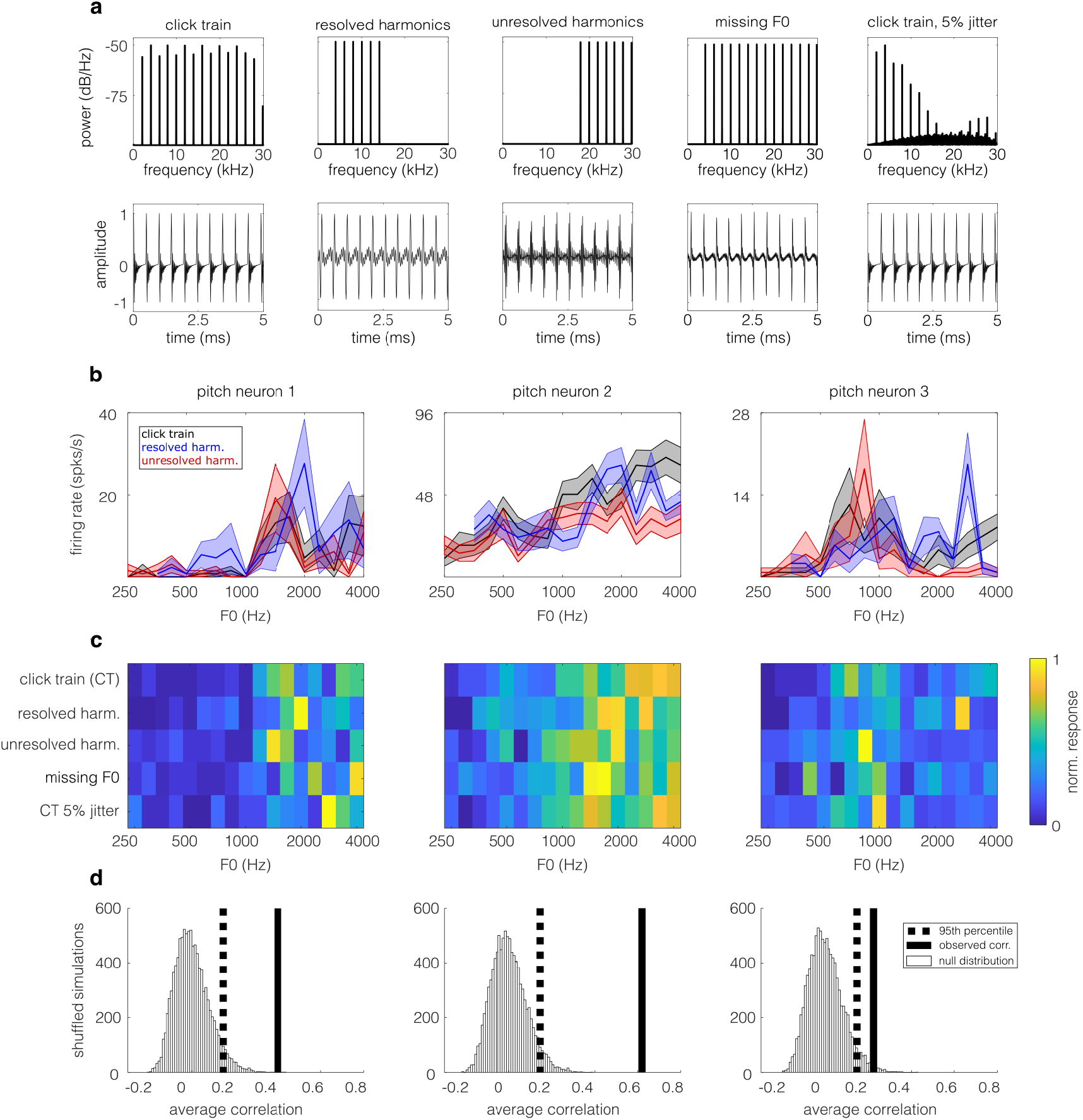
Pitch neurons can derive F0 from resolved and unresolved harmonics. a) Spectra (top row) and waveforms (bottom row) of 5 different sound types used to classify pitch neurons (F0 = 2000 Hz). b) Trial-averaged spike rate responses as a function of F0 for three example pitch neurons. Pitch neurons can derive F0 from resolved or unresolved harmonics and therefore have similar F0-tuning to click trains (black line), resolved harmonics (blue line), and unresolved harmonics (red line). c) Normalized trial-averaged responses to 5 types of stimuli that contain resolved or unresolved harmonics, or both, plotted as in Fig. 1c. d) Results of bootstrapped correlation test of F0-tuning similarity across stimuli, plotted as in Fig. 1d. The same 3 example temporal neurons are shown in b, c and d.

Our bootstrapped correlation analysis showed that 31 of the 32 putative pitch neurons had statistically similar F0-tuning across all 5 stimuli (p<0.05; Fig. 4d), even when we include sounds with non-overlapping frequency spectral (namely, low and high harmonic stimuli). This evidence extends the existence of acoustically invariant “pitch neurons” beyond marmosets^21^, revealing a conserved cortical mechanism for pitch coding.

### Sensitivity to pitch salience

If harmonicity and temporal neurons contribute to ferrets’ percept of pitch, we might expect their response magnitude to decrease as pitch salience decreases. In human listeners^22,23^, pitch salience decreases as the temporal regularity of a click train is degraded by randomly “jittering” the timing of individual clicks. Pitch neurons in the marmoset have been shown to be sensitive to this jittering of click trains^21^. We investigated if pitch selective neurons in ferrets are modulated by pitch salience, by presenting click trains with 5 levels of temporal jitter (0%, 5%, 10%, 20%, and 40% jitter; Fig. 5a).

**Figure 5.**
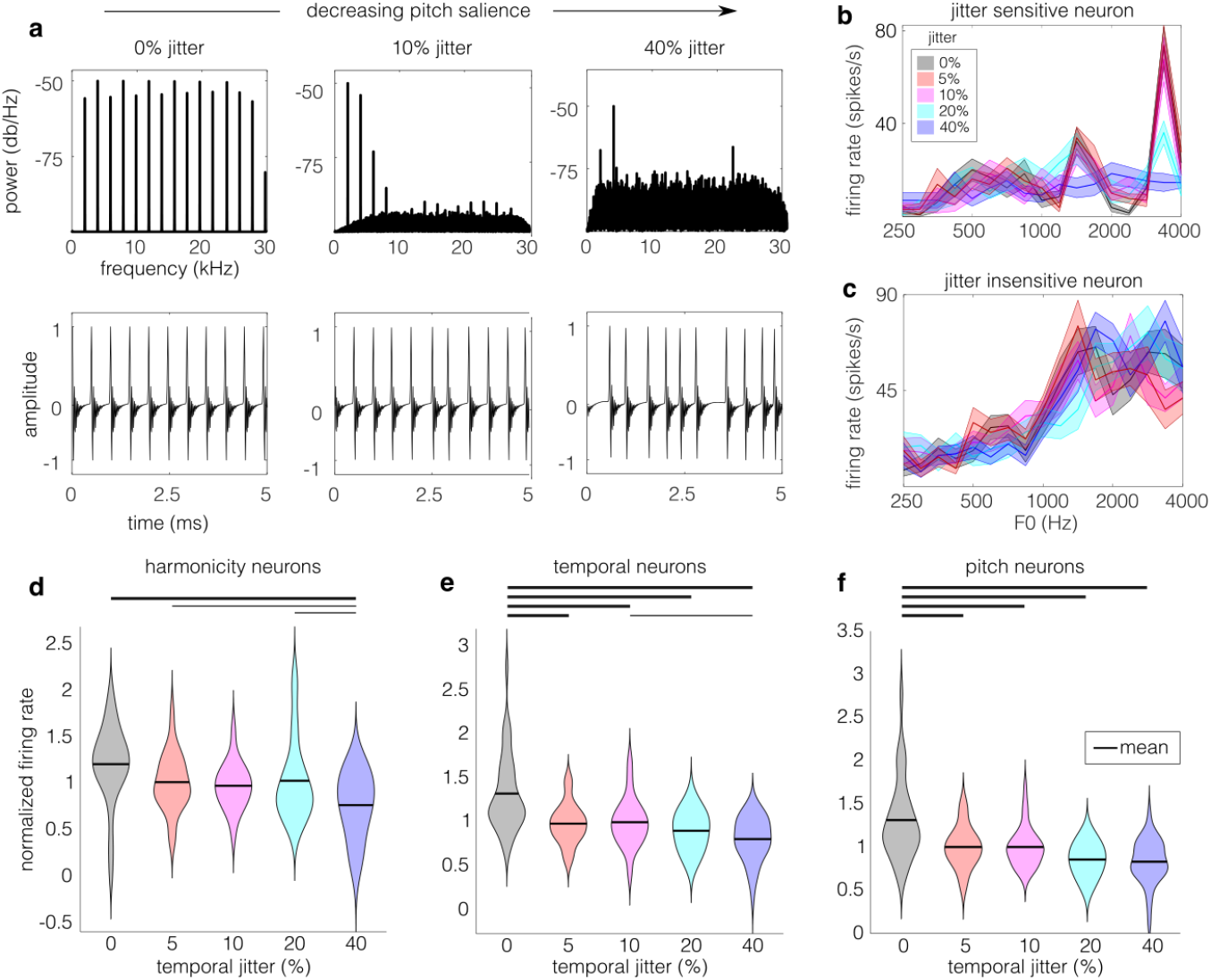
Temporal and harmonicity neurons are sensitive to temporal regularity. **a**) Spectra (top row) and waveform (bottom row) of click trains with increasing temporal jitter between clicks (left to right). Jittering the temporal regularity of the clicks in the click train introduces noise in the harmonic spectrum (top row) and periodicity (bottom row), reducing the salience of the perceived pitch in human listeners^22,23^. **b**) F0-tuning of an example harmonicity neuron in response to click train stimuli with 0, 5, 10, 20, or 40% temporal jitter. The tuning curve is consistent for low percentages of jitter, but becomes flat for jitter above 20%. **c**) F0-tuning for an example temporal neuron, plotted as in **b.**The F0-tuning of this neuron is not sensitive to the jitter manipulation. **d**) For each harmonicity neuron, the normalized response at best F0 (+/-0.25 octaves) was calculated for each jitter level. The responses of neurons decreased as jitter level increased (1-way ANOVA with Tukey-Kramer HSD test; thin line: p<0.05, thick line: p<0.001). **e**) Jitter sensitivity at best F0 for temporal neurons, plotted as in **d. f**) Jitter sensitivity at best F0 for pitch neurons, plotted as in **d**.

An example jitter-sensitive harmonicity neuron is shown in Fig. 5b. Its spike rate at best F0 decreases as jitter increases. For comparison, Fig. 5c shows a temporal neuron which is not sensitive to jitter; its tuning curve and response magnitude are consistent across jitter levels. Therefore, some neurons classified as harmonic and temporal neurons were sensitive to this manipulation of temporal regularity. Next, we examined if the populations of harmonicity and temporal neurons were jitter-sensitive. The spike rate response of each neuron was calculated for click trains presented within a half octave of best F0, and for each of the 5 levels of jitter. Responses were normalized by the average response across all jitters presented to remove variance in overall firing levels across neurons. We found that harmonicity (1-way ANOVA; F(4)=6.24, p=1×10^-4^; Fig. 5d), temporal (F(4)=16.5, p=1.6×10^-11^; Fig. 5e), and pitch (F(4)=13.6, p=1.8×10^-9^; Fig. 5d) neurons all showed weaker spiking responses as the level of jitter increased. Therefore, all three classes of pitch selective neurons encode temporal regularity in a manner consistent with a neural representation of pitch salience.

### F0-tuning of pitch selective neurons is not predicted by pure tone frequency tuning

The above analyses have shown that a subpopulation of auditory cortical neurons can derive the fundamental frequency that we hear as pitch from resolved harmonics (harmonicity neurons), temporal periodicity (temporal neurons), or both these types of acoustical cues (pitch neurons). While pure tones elicit a percept of pitch in humans and ferrets^24^, they do not contain harmonics from which to derive harmonic relations, and they only provide temporal periodicity cues in one frequency channel of the labelled line. Thus, they would not provide strong inputs for the type of harmonic and temporal neurons we have proposed here, which integrate across frequency channels. Previous studies have shown a poor correspondence between pure tone frequency tuning and preferences for the F0 of artificial vowel sounds in auditory cortex^25^, and pure tones are also poor predictors of cortical responses to complex sounds more generally^26^. Therefore, we hypothesize that pure tone responses may be a poor predictor of F0-tuning for complex sounds in the harmonic, temporal, and pitch neurons described here.

We tested this hypothesis by comparing the F0-tuning of harmonicity, temporal, and pitch neurons measured in response to click trains (containing resolved harmonic and temporal periodicity cues), low harmonic complexes (resolved harmonics only), high harmonic complexes (temporal periodicity without resolved harmonics), and pure tones (energy at F0 only). Fig. 6a shows the tuning curves for these 4 stimuli for 3 example pitch neurons. The first example neuron (green border) shows considerable overlap in its pure tone frequency preference (double-peaked tuning for 1414 and 2828 Hz) and best F0 for complex sounds (F0 preference for one of these two frequencies for each complex sound). However, the other two example neurons display consistent F0 preferences for the 3 complex sounds, but non-overlapping frequency tuning derived from responses to pure tones.

Overall, our harmonicity, temporal, and pitch neurons showed F0-tuning to complex sounds that did not readily correspond to their pure tone frequency tuning. The scatterplot in Fig. 6b shows the correlation of F0-tuning in pitch neurons between the click trains and resolved harmonic stimuli (x-axis) and between the click trains and unresolved harmonic stimuli (y-axis). The clustering of neurons in the top right quadrant indicates that their F0-tuning measured with broadband click trains was similar that that measured with both resolved and unresolved harmonics (Fig. 6b; resolved harmonic correlations: µ=0.46, σ=0.26; unresolved harmonic correlations: µ=0.35, σ=0.32). This structure was not observed for pure tone stimuli (Fig. 6c; resolved harmonic correlations: µ=0.18, σ=0.31; unresolved harmonic correlations: µ=0.06, σ=0.28). Similarly, the F0-tuning of harmonic (µ=0.02, σ=0.39) and temporal neurons (µ=0.15, σ=0.30) were not positively correlated with their pure tone frequency tuning (Fig. 6b, marginal histograms). These results demonstrate that complex pitch tuning in these neurons is distinct from the mechanisms used to encode a single pure tone frequency.

**Figure 6.**
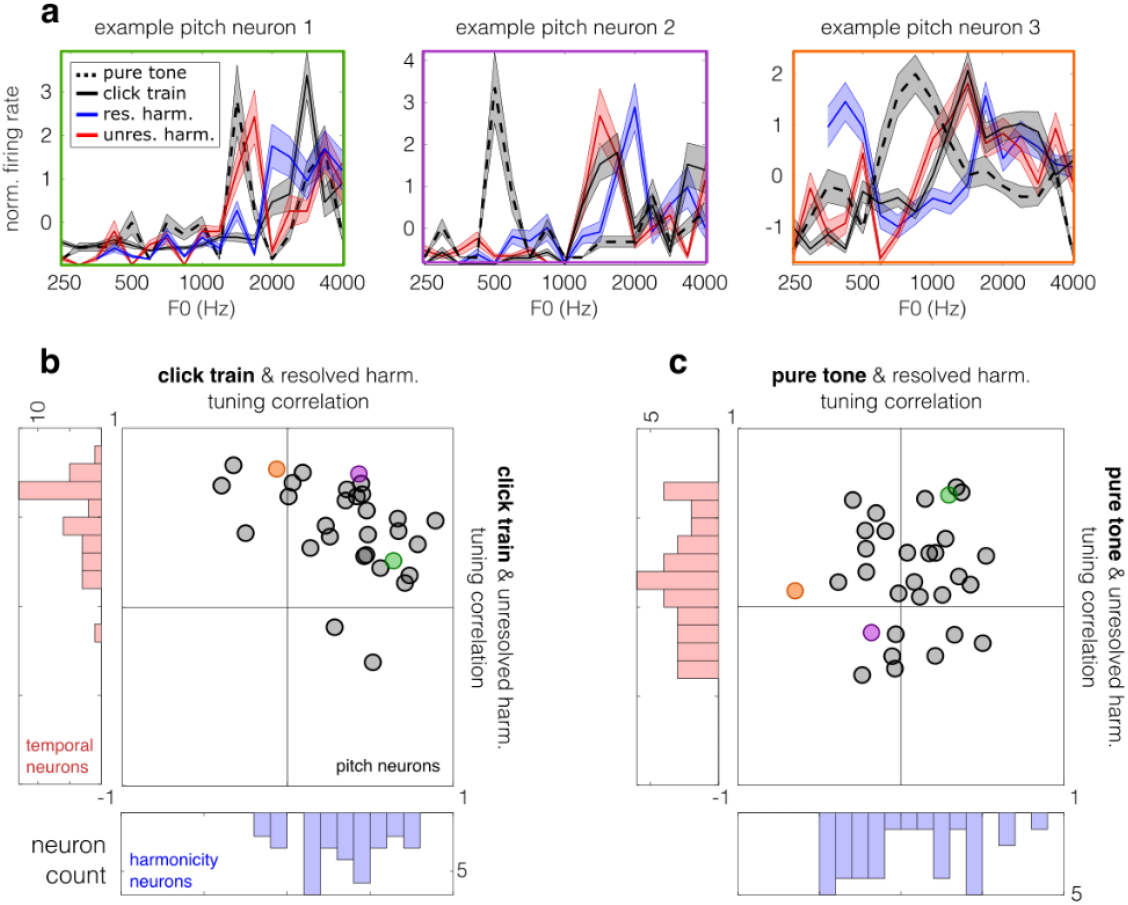
Pitch neurons’ F0-tuning for complex sounds does not correspond to pure tone frequency tuning. **a**) Pure tone frequency tuning and F0 tuning curves are shown for three 3 pitch neurons. Normalized trial-averaged spike rates (lines) are shown with SEM (shaded regions). The green, purple, and orange borders around these examples indicate their identity in **b** and **c. b**) Axes indicate the correlation between F0-tuning measured with click trains and low harmonics (x-axis) or high harmonics (y-axis). F0-tuning to click trains is positively correlated with F0-tuning to low harmonics in nearly all harmonicity neurons (blue histogram), and it is positively correlated with F0-tuning to high harmonics in nearly all temporal neurons (red histogram). F0-tuning to click trains is positively correlated with F0-tuning for both low and high harmonics in most pitch neurons (scatter plot; each dot is one pitch neuron). **c**) Results plotted as in **b**, but examining correlations with pure tone frequency tuning, rather than click train F0-tuning. The distribution of correlations between frequency tuning and F0-tuning to low harmonics is centered around zero in harmonicity neurons (blue histogram). Similarly, temporal neurons’ frequency tuning is not correlated with F0-tuning to high harmonics (red histogram). In pitch neurons, pure tone frequency tuning does not correlate systematically with F0-tuning to low or high harmonics (scatter plot).

### Cochlear distortion products cannot account for the observed F0-tuning

When a missing fundamental sound is presented to the ear, the active mechanism of outer hair cell amplification causes displacement of the basilar membrane at the position corresponding to F0^27,28^. Perceptually, psychophysical studies have shown that cochlear distortion products can improve pitch detection in human subjects^29,30^. Therefore, it is important to determine if cochlear distortion products contribute to the F0-tuning we observed in cortical neurons in response to our missing fundamental stimuli, particularly our high harmonic stimuli which were designed to only contain unresolved harmonics. Distortion products at F0 are 10-15 dB quieter than the harmonic components in a missing fundamental sound^29^, so we presented our low and high harmonic stimuli in the presence of pink noise (5 dB below the harmonic components) to mask potential distortion products in the ferret cochlea. If the observed F0-tuning persists in these neurons even in the presence of the pink noise masker, this tuning is not entirely produced by distortion products.

We found that pitch selective neurons retained their F0-tuning even in the presence of a pink noise masker. Fig. 7a,c show similar F0-tuning with and without masker for example harmonicity and temporal neurons. This finding held for the population of harmonicity and temporal neurons. F0-tuning curves for masked and unmasked resolved harmonics were more similar for harmonicity neurons than other neurons (two sample t-test; t(839)=8.7, p=2.5×10^-17^; Fig. 7b). Similarly, the F0-tuning for unresolved harmonics presented with and without a pink noise masker were significantly similar in temporal neurons (t(843)=9.8, p=2.2×10^-21^; Fig. 7d). Furthermore, the preferred F0 for masked and unmasked versions of the sounds were significantly correlated in harmonicity and temporal neurons (Pearson correlation; r=0.55, p=8.7×10^-4^ for harmonicity neurons; r=0.57, p=2.0×10^-4^ for temporal neurons; Fig. 7e,f). These results confirm that the pitch selectivity observed here is not due to cochlear distortion products.

**Figure 7.**
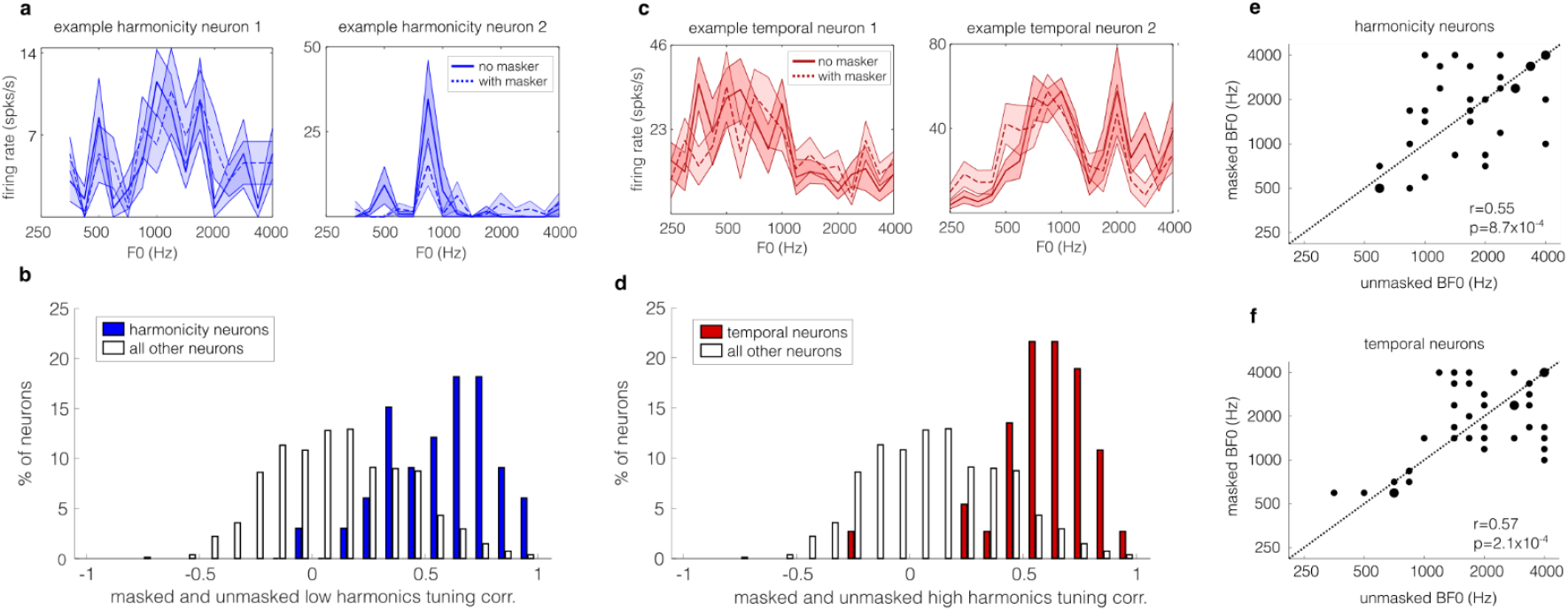
Distortion products do not explain observed F0-tuning. **a**) Each panel shows data from one example harmonicity neuron. Trial-averaged spike rates are plotted as a function of F0 for low harmonics presented with (dashed line) or without (solid line) a pink noise masker. **b**) Histograms show that the correlation between F0-tuning for masked and unmasked low harmonics within harmonicity neurons (blue bars) was higher than in other sound-responsive neurons (open bars). **c**) Each plot shows F0-tuning for masked (dashed line) and unmasked (solid line) high harmonics for two example temporal neurons, plotted as in **a. d**) Plotted as in **b**, histograms show the correlation between F0-tuning for masked and unmasked high harmonics within temporal neurons (red bars) and other sound-responsive neurons (open bars). **e**) For each harmonicity neuron, the scatter plot shows the best F0 measured with unmasked and masked low harmonics. Larger dots indicate two harmonicity neurons. **f**) Best F0 of temporal neurons for unmasked and masked high harmonics, plotted as in **e**.

### Anatomical locations of harmonic, temporal and pitch neurons

Given the evidence for an anatomically segregated region of pitch neurons in the marmoset auditory cortex^21^, we next explored where harmonicity, temporal, and pitch neurons were located across the surface of ferret auditory cortex. The auditory cortex of each animal was mapped based on anatomical landmarks (e.g. sulci), latency of neural responses to tones and noise bursts, and best frequencies (as described previously)^18^. Using this mapping, each neuron was categorized to low- or high-frequency regions of A1 or AAF. The proportion of F0-sensitive neurons classified as harmonicity, temporal, or pitch neurons is mapped in each region is summarized in Supplementary Table 1 and visualized in Supplementary Fig. 1. Only 1 F0-sensitive neuron was located in low-frequency AAF, so we omitted this region from further analysis.

To assess whether the three populations of neurons were homogeneously distributed across primary auditory cortex, we applied a chi-square test for homogeneity, with the anatomical regions as the categorical variable. We found that harmonicity neurons were distributed similarly across the three cortical regions (low frequency A1, high frequency A1, and high frequency AAF; χ^2^(2)=1.78, p=0.41; Fig. 8a). Temporal neurons (Fig. 8b) and pitch neurons (Fig. 8c) were significantly more prevalent in high-frequency A1 and AAF, particularly high-frequency AAF (χ^2^(2)=25.6, p=2.7×10^-6^ for temporal neurons; χ^2^(2)=16.6, p=2.5×10^-4^ for pitch neurons). In summary, we did not find strong evidence for a single pitch centre in auditory cortex, although the proportion of temporal and pitch neurons varied between primary fields and across the tonotopic gradient.

**Figure 8.**
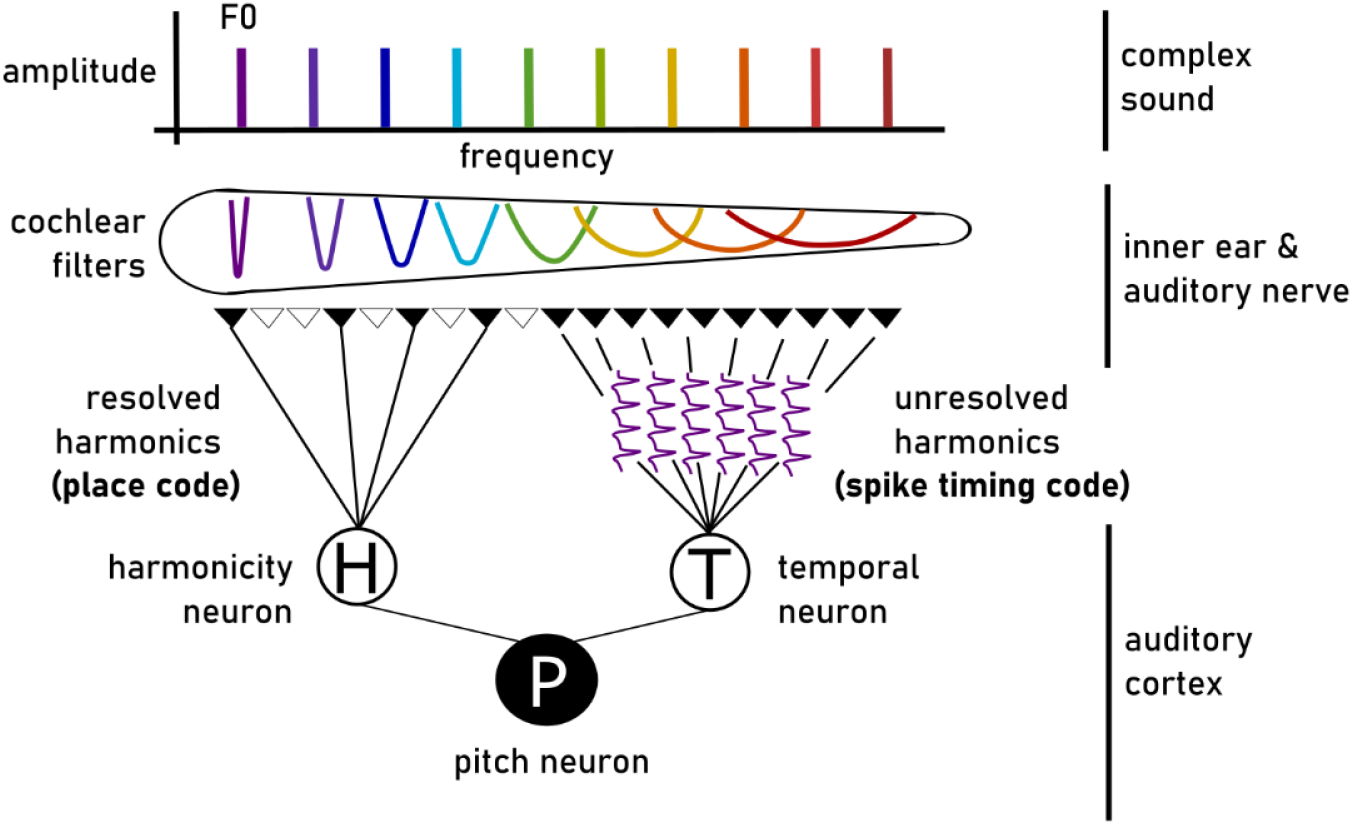
Schematic of proposed dual extraction of fundamental frequency (F0) from resolved and unresolved harmonics. Top: The spectrum of a harmonic tone complex is shown, with each frequency component in a different color. Middle: The frequency tuning curves of individual cochlear filters are shown along the unrolled basilar membrane, with color indicating the best frequency of each auditory nerve fiber. Because the tuning bandwidth of auditory nerve fibers is constant with respect to logarithmic frequency, tuning curves become exponentially broader with increasing linear frequency. Low numbered harmonics are termed “resolved” if only one harmonic falls within the tuning curve of a single auditory nerve fiber, and therefore F0 is represented with a place code as the pattern of harmonic activation across the population of resolved fibers. In contrast, auditory nerve fibers tuned to higher harmonics are likely to respond to multiple harmonics in the sound, so these are “unresolved” in the place code. Because multiple unresolved harmonics excite the same auditory nerve fiber, the spike times of auditory nerve fibers can phase lock to the sum of unresolved harmonics at F0 (purple traces show spike output over time). This produces an explicit temporal representation of F0 for resolved harmonics. Bottom: Our data show that there are distinct neural populations in auditory cortex to extract these F0 cues: harmonicity neurons (H) compute F0 from the place code of exclusively resolved harmonics; and temporal neurons (T) extract F0 from the temporal code of unresolved harmonics. These two neural populations may then converge on a third neural population, pitch neurons (P), which encode F0 irrespective of the types of acoustic cues present in the sound.

## Discussion

The fundamental frequency (F0) of harmonic sounds can be computed from either harmonic spacing or temporal periodicity^31,32^. Computational models have shown that a single spectrotemporal algorithm for F0 extraction is also plausible^33,34^, but human psychophysical studies are not consistent with such an algorithm, and instead suggest that pitch arises from the parallel processing of harmonic and temporal cues^9,35^. Behavioral studies have demonstrated that ferrets, like humans, can discriminate pitch shifts in complex sounds^24,36^, although ferrets base their pitch judgments more heavily on temporal periodicity than resolved harmonic cues^12^. Previous studies have shown that neurons throughout ferret auditory cortex can represent the F0 of artificial vowels, and almost always in parallel with other sound features^18,25,37^, and that their activity correlates with pitch judgments during behaviour^38^. However, it is not clear if neurons in ferret auditory cortex represent F0 more generally across stimuli, process harmonic and temporal cues to extract F0, nor if they include specialized pitch neurons.

Here, we used high-density microelectrode recordings and a diverse stimulus set to identify three new neural subpopulations in ferret primary auditory cortex: harmonicity neurons that encode the F0 of resolved harmonics; temporal neurons tuned to the periodicity of F0 in unresolved harmonics; and pitch neurons that can extract F0 from both cue types. These populations showed properties previously described in primate pitch neurons, including sensitivity to phase manipulations and temporal regularity, responses to missing fundamentals at F0, and invariance to cochlear distortion products^21,39,40^. These F0 computations are well supported by findings of complex pitch discrimination in behaving ferrets^12,24^, and could allow a listener to generalize pitch across a wide array of sounds, as humans do when recognizing a melody across different instruments.

A model of how harmonicity, temporal and pitch neurons may extract F0 from the representations of complex sounds in the cochlea is presented in Fig. 8. Harmonicity neurons (H) could integrate inputs across labelled line representations of the resolved harmonic components of a sound. This mechanism has been previously described as a “harmonic sieve”, with excitatory inputs from neurons tuned to individual harmonics of the preferred F0 and inhibitory inputs from intermediate frequencies^41^. Temporal neurons (T) may instead involve circuits that compute the periodicity of spike times in neurons tuned to unresolved harmonics. These inputs may also include neurons phase-locked to resolved harmonic components, as F0 can in theory be calculated as the autocorrelation of spike times across neurons tuned to different harmonics^42^. Pitch neurons (P) can provide spectrally-invariant representations of F0, perhaps by combining inputs from harmonic and temporal neurons tuned to a common F0. The auditory nerve representations of resolved harmonic and temporal periodicity described in Fig. 8 have been demonstrated in the microelectrode studies of Delgutte and colleagues^8^. The evidence for the proposed dual processing of pitch cues at the level of auditory cortex is summarized below.

### Dual pitch processing in auditory cortex

Our findings add to the evidence in non-human primates that cortical neurons can represent F0 using resolved harmonic or unresolved temporal envelope cues^10,40,43^. In macaques, populations of neurons in low-frequency A1 can represent resolved harmonics as a place code, while higher-frequency neurons time-lock to the temporal envelopes of unresolved harmonics^20,43^. In marmosets, Wang and colleagues identified a “pitch area” on the low-frequency border of A1 and R, where individual neurons were tuned to the F0 of both pure tones and missing fundamentals^21,39^.

Within the marmoset pitch centre, a small subpopulation of pitch neurons were sensitivity to alternating phase manipulations^40^. A separate subset of neurons with higher best F0s (>450 Hz) was insensitive to phase manipulations, but sensitive to resolved harmonic cues^40^. Ferret auditory cortex shows a similar division of labour. Ferret temporal neurons show sensitivity to phase manipulations, including a reduced response to both alternating phase and random phase harmonics (Fig. 3). They show similar tuning across complex periodic stimuli, even when only unresolved harmonics are presented (Fig. 2). Harmonicity neurons in our study generalized their F0-tuning across complex stimuli, but only for sounds containing resolved harmonics (Fig. 1). As reported for marmoset pitch-selective neurons^21,39^, our temporal and harmonicity neurons were sensitive to the temporal regularity of click trains (Fig. 5), demonstrating a neural parallel to pitch salience in human listeners^22,23^. Our results suggest that dual cortical processes for extracting F0 may be widespread across mammals.

### Pitch-selective neurons in auditory cortex

Our results demonstrate that pitch-selective neurons exist in ferrets. As reported in marmosets, some pitch-selective neurons were sensitive to the temporal regularity of jittered click trains, were unaffected by the presence of a masking noise to control for cochlear distortion products, and were sensitive to the phase of unresolved harmonic components^21,39^. Their rarity was also comparable: about 3.5% of sound-responsive neurons met our pitch neuron criteria, closely matching the ∼3.6% reported in marmoset auditory cortex (Fig. 2 of Bendor and Wang, 2005^21^), although about one-third of neurons are pitch selective in the marmoset pitch center^21,39,40^. To our knowledge, this is the first demonstration of pitch-selective neurons outside primates.

Two differences between marmoset and ferret pitch neurons are noteworthy. First, ferret pitch neurons were not typically well driven by pure tones, while marmoset pitch neurons were tuned to similar F0s for pure tones and missing fundamentals^21,40^. This may reflect our differences in classification criteria, but it also raises the possibility that pure-tone frequency and complex F0 extraction are mediated by distinct mechanisms. A single tone has limited harmonic structure, providing a weak input to our model of temporal and harmonicity neurons, which both pool information across harmonics (Fig. 8).

A second difference is that temporal, harmonicity, and pitch neurons were found distributed throughout low and high frequency primary auditory fields in ferrets, albeit in variable proportions. This contrasts with the local pitch area of marmosets^21^, but aligns with more distributed pitch representations reported in auditory cortex of macaques^20,43,44^ and humans^45–48^. Even human fMRI studies showing evidence of a pitch center disagree on its location within auditory cortex^44,49,50^. Overall, the bulk of evidence, suggests that a distributed network of pitch-selective neurons may be the more general organizational principle. Anatomical clustering in marmosets may represent a species-specific specialization, or could reflect the criterion for pitch neurons to generalize their F0 tuning to pure tones.

## Conclusions and future directions

These findings establish that ferret auditory cortex implements dual pitch extraction strategies and contains neurons specialized for invariant pitch representation. This supports the idea that pitch perception relies on conserved cortical computations across mammals, including non-primates, and opens the door to studies of pitch in a wider range of species. For example, if mice were shown to experience pitch perception, a toolbox of power genetic manipulations would be available to further dissect pitch processing circuits.

Future work should test the causal role of these neurons in behavior. Despite the wealth of evidence for anatomically localized pitch neurons in the marmoset, the behavioural consequences of manipulating this area have not been explored. Comparative studies should also clarify how differences in cortical organization relate to species’ ecological or trained demands. For example, do species with cochlea that provide fewer resolved harmonics (e.g. mice) naturally have more neurons tuned to temporal than harmonic pitch cues? Would training on resolved harmonics result in a higher proportion of temporal pitch neurons? Many key questions about the neural mechanisms of pitch perception remain, and the identification of ferret pitch selective neurons suggests that these questions can be answered in a wide range of animal models.

## Methods

### Animals

Two female and two male adult ferrets (age: µ = 29.3 weeks, σ = 18.4 weeks; *Mustela putorius furo*; Marshall BioResources, UK) were used in this study. The animal procedures in this study were approved by the local ethical review committee of the University of Oxford and were performed under UK Home Office license.

### Surgical Procedure

We performed terminal electrophysiological recordings on each ferret. The animals were put under general anesthesia via an intramuscular injection of ketamine (Vetalar; 5 mg/kg) and medetomidine (Domitor; 0.02 mg/kg). Anesthesia was maintained for the duration of the surgery and electrophysiology with a continuous intravenous infusion of ketamine and medetomidine in Hartmann’s solution with 3.5% glucose and dexamethasone (0.5 mg/ml/hr) via intravenous cannula. The ferrets were intubated and artificially ventilated with medical O_2_. Blood oxygenation, electrocardiogram, respiratory rate, and end-tidal CO_2_ were continuously monitored throughout the recording session, and a body temperature of 36-38°C was maintained with a heating pad and warm air (3M Bair Hugger). To prevent corneal desiccation, eye ointment (Maxitrol; Alcon, UK) was applied throughout the procedure. Every 6 hours, the animals received atropine (Atrocare; 0.06 mg/kg i.m.), or when bradycardia or arrhythmia was observed.

Animals were placed in a custom stereotaxic frame and secured in place with a mouthpiece and ear bars. The scalp was then shaved, cleaned (ChloraPrep; 2% chlorhexidine gluconate), and injected with bupivacaine (Marcain, <1 mg/kg s.c.). The scalp was then incised, and the temporal muscles retracted. Using dental cement (SuperBond; C&B, UK) and a stainless steel bone screw (Veterinary Instrumentation, UK), a steel holding bar was attached to the skull. A craniotomy (10 mm diameter) was carried out over the right auditory cortex, and the exposed dura was removed. Anatomical location was confirmed from stereotaxic coordinates (11 mm ventral to the midline and 8 mm anterior to Lambda) and visual identification of the ectosylvian gyrus. The surface of the brain was covered with a solution containing 1.25% agarose in 0.9% NaCl. Regularly throughout recording, silicone oil was applied to the craniotomy.

Immediately prior to recording, the ear bars were removed, and the animal with the frame was transferred to an electrically isolated anechoic chamber. An Ag/AgCl reference wire was inserted between the skull and the dura at the edge of the craniotomy. A microelectrode probe (Neuropixels Phase 3)^51^ was inserted through the entire depth of auditory cortex, orthogonally to the brain surface. The cortical area probed by each penetration was identified based on its location relative to the ectosylvian gyrus, the local field potential response latencies, and the frequency response area (FRA) shapes of cortical units recorded. Across the 18 cortical penetrations that yielded sound-responsive units, 10 were in low frequency A1, 3 in high frequency A1, 2 in low frequency AAF, and 3 in high frequency AAF. After a complete presentation of the stimulus set, the probe was removed and reinserted in a new location within auditory cortex. Data were acquired using the SpikeGLX software (https://github.com/billkarsh/SpikeGLX)^52^ and custom MATLAB scripts (Mathworks) at a 30 kHz sampling rate.

### Sound presentation

Stimuli were presented binaurally via earphones (Panasonic RP-HV094E-K) driven by a System 3 RP2.1 multiprocessor and headphone amplifier (Tucker-David Technologies). The earphones were closed-field calibrated using a GRAS amplifier and 1/8 inch condenser microphone placed at the end of a model ferret ear canal. Inverse filters were designed to ensure that the devices produced flat (less than ±3 dB) outputs across 200 Hz – 30 kHz. During the experiment, the earphones were coupled with otoscope speculae which were inserted into each ear canal and sealed in place with Otoform (Dreve Otoplastik). Sounds were presented at 48,828 Hz sampling rate.

### Sound stimuli

The main stimulus set consisted of 13 sound types: pure tones and 12 different complex sounds, each presented over a range of 17 F0s (200-4000 Hz in 0.5 octave increments). There were three broad categories of acoustic manipulations in our complex stimuli: temporal regularity, harmonic components, and phase. In the temporal regularity category, we presented click trains that were composed of biphasic pulses at a rate of F0. The 5 different types of “click train” stimuli were composed with different levels of temporal jitter between clicks (0, 5, 10, 20, and 40%). In the 0% jitter condition, the clicks were perfectly periodic, occurring with a spacing of 1/F0. In the jittered condition, each click from a regular click train was randomly shifted in time. The maximal amount of time shift was defined as a percentage of the inter-click interval (5, 10, 20, or 40%), centered on the timing of that click in a regular click train^21,22^. Therefore, stimuli with an increased jitter are less temporally regular, and these are perceived by humans^22^ and ferrets (unpublished data from our lab) to have a weaker pitch salience.

The second category, harmonic component manipulations, comprised harmonic sounds designed to include either: 1) energy at all harmonics within our passband except at F0 (200 Hz – 30 kHz; “missing F0”); 2) only harmonics that would be resolvable on the ferret cochlea (“low harm”); or 3) only harmonics that are unresolved for ferrets (“high harm”). The resolved frequencies for each F0 were determined using a model of ferret cochlear filters based on ferret auditory nerve recordings^12,53^. In addition, resolved and unresolved harmonic stimuli were presented with and without a pink noise masker (“low mask” and “high mask”) to mask cochlear distortion artefacts. The masker was presented at a sound level that was 5 dB below the harmonic complex tone. In this condition, the noise masker would be 10 to 15 dB above the level of the expected individual tonal components at F0^29^.

For the phase manipulations, the third category, we presented unresolved harmonic stimuli with either randomized phases across all harmonics (“rand. phase”), or with neighboring harmonics in alternating opposite phases (sine and cosine phase, “alt. phase”). These stimuli were generated by adding sin waves at the frequency of the desired harmonics for each combination of F0 and harmonic content. These stimuli did not include energy at F0. All sounds were generated with custom Matlab scripts, and bandpass filtered from 200 Hz – 30 kHz.

Each sound was presented for either 200 ms (3 ferrets) or 300 ms (1 ferret) at 70 dB SPL_A_. Each unique combination of sound type and F0 (e.g. resolved harmonics at 1000 Hz) was presented 13 times, in pseudo random order across type and F0, with an interstimulus silent interval of 600 ms (3 ferrets) or 400 ms (1 ferret), and linear onset and offset ramps of 25 ms.

In addition, a fuller set of pure tone stimuli was presented to assess frequency tuning in each penetration, so that we could localize each of our microelectrode penetrations on the tonotopic map. Broadband gaussian noise bursts of 100 ms were used as a search stimulus to identify sound-responsive neurons at the beginning of each electrode penetration, followed by presentation of pure tones for frequency tuning classification, and then the full set of pitch stimuli described above. Pure tones were presented at one of 15 different frequencies (500 Hz – 40 kHz in 0.45 octave increments) at 40, 50, 60, 70, or 80 dB. Tones lasted 100 ms followed by 500 ms of silence. Each unique combination of tone frequency and intensity was presented 20 times to capture the frequency response area of the electrode shank. FRAs were recorded for each penetration to identify the tonotopic gradients across AAF and A1.

### Spike sorting

The recorded signal was processed offline by first digitally high-pass filtering at 150 Hz. Common average referencing was performed to remove noise across electrode channels^54^. Spiking activity was then detected and clustered using Kilosort2 software (https://github.com/MouseLand/Kilosort2)^55,56^. Responses from single units were manually curated using Phy (https://github.com/cortex-lab/phy)^57^. Units showing stereotypical spike shapes with low variance, and a clear refractory period in their autocorrelation spike histogram were classified as single units. Only well isolated single units were included in subsequent analyses.

### Isolating F0-sensitive neurons

Each neuron was first assessed for sound responsivity using a paired-sample t-test (alpha = 0.05), which compared spike rates during 150 ms of silence immediately prior to sound onset to rates during the 150 ms after sound onset across all presentations of our sound stimuli. Only neurons found to be significantly responsive to sound were included in subsequent analyses. We assessed neural sensitivity to F0s in response to each of the 13 sound types using a one-way ANOVA (alpha = 0.05), with spike rates during the first 100 ms after sound onset as the dependent variable and F0 as the grouping variable. The best F0 for a given neuron was taken as the F0 eliciting the largest trial-averaged spike rate for a given stimulus type (e.g. click trains).

### Assessing tuning similarity

We assessed the similarity of a neuron’s F0-tuning across pairs of sound types using a bootstrapped sampling procedure. First, F0 tuning curves were constructed for a given neuron and for each stimulus type by averaging the spike rate in the first 100 ms after sound onset across trials. Next, we calculated the correlation of F0-tuning curves for every possible pair of sound types for that neuron. These pairwise correlations were averaged across stimulus pairs to create a single metric of how consistent the neuron’s F0-tuning was across different sound types. To test whether this average correlation of F0-tuning was greater than would be expected by chance, we created a null correlation distribution from “shuffled” tuning curves, in which the neuron’s average responses were randomly re-assigned across F0s. We repeated this procedure for 10,000 shuffled versions of each neuron’s dataset. If the pairwise correlation of the unshuffled tuning curves was greater than the 95^th^ percentile of the null distribution, the neuron was considered to have consistent F0-tuning across the different sound types.

### Declaration of generative AI and AI-assisted technologies in the manuscript preparation process

During the preparation of this work the authors used ChatGPT in order to make suggestions for reducing the abstract word count. After using this tool, the authors reviewed and edited the content as needed and take full responsibility for the content of the published article.

## Author contributions

VT analyzed the data and wrote the manuscript. QG collected the data, analyzed the data, and contributed to the manuscript. KMMW conceptualized the experiment, collected the data, edited the manuscript, and secured the funding.

## Acknowledgements

This work was funded by Biotechnology and Biological Sciences Research Council grants to KMMW (BB/M010929/1, BB/X013103/1) and a University of Oxford Clarendon Scholarship to VT. We are grateful to Dr Ben Willmore and Dr Aleksandar Ivanov for assisting with data collection, to Prof Randy Bruno for helpful comments on a draft manuscript, and to Prof Andrew King for sharing experimental equipment for this study.

