## Supplementary Figures for "Pitch selectivity in ferret auditory cortex"

| cortical region | penetrations | sound-responsive neurons | F0-sensitive neurons | harmonicity neurons | temporal neurons | pitch neurons |
|---|---|---|---|---|---|---|
| low-frequency A1 | 10 | 566 | 103 | 20 | 11 | 11 |
| high-frequency A1 | 3 | 180 | 28 | 3 | 9 | 6 |
| low-frequency AAF | 2 | 33 | 1 | 0 | 0 | 0 |
| high-frequency AAF | 3 | 93 | 33 | 4 | 17 | 14 |

Supplementary Table 1. Distribution of cortical penetrations across the four primary auditory cortical regions probed. We report the total number of sound-sensitive, F0-sensitive, harmonicity, temporal, and pitch neurons across all penetrations for each region.

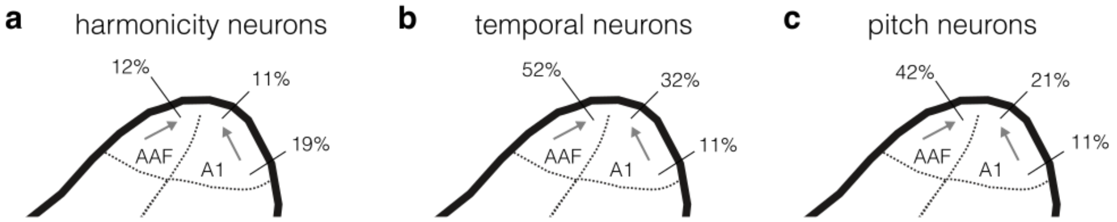


Supplementary Figure 1: Distribution of harmonicity, temporal, and pitch neurons across primary auditory cortex. **a**) Percentages indicate the proportion of F0-sensitive neurons that were also classified as harmonicity neurons. Arrows indicate low-to-high frequency tonotopic gradient. **b**) Distribution of temporal neurons, plotted as in **a**. **c**) Distribution of pitch neurons, plotted as in **a**.
